# Type I collagen gels for assessing the combined effects of ligand concentration-dependent elasticity and fibril density on cells

**DOI:** 10.1101/2025.03.10.642521

**Authors:** Tomoko Gowa Oyama, Kotaro Oyama, Hiromi Miyoshi, Mitsumasa Taguchi

## Abstract

Type I collagen (Col-I) is the most abundant fibrous protein in the extracellular matrix (ECM); its densification is involved in conditions including cancer, fibrosis, vascular diseases, and aging. Increased Col-I concentrations lead to increased ECM fibril density and stiffening; therefore, the effects of such changes in the microenvironment of cells require examination. Three types of Col-I gels were developed by introducing reagent-free radiation-induced crosslinking, while adjusting the distance between the Col-I monomers or fibrils. The adhesion area of HeLa human cervical cancer epithelial cells, human induced pluripotent stem (hiPS) cells, and 3T3-Swiss albino mouse embryonic fibroblasts, tended to decrease with increasing fibril density when the ligand concentration-dependent elasticity (*C*-*E*) was constant. In the absence of fibrils, the adhesion area of HeLa and 3T3-swiss cells increased with increasing *C*-*E*, whereas hiPS cells clustered regardless of *C*-*E*. When *C*-*E* and the fibril density increased simultaneously, the adhesion area of HeLa and 3T3-swiss cells remained unchanged due to the two opposing effects. Our results highlight the importance of evaluating the combined effects of the compositional, mechanical, and topographical cues of Col-I on cells. The developed Col-I gels will contribute to elucidating the role of Col-I in development, differentiation, regeneration, disease, and aging.

**Highlights:**

- Col-I concentration-dependent elasticity (*C*-*E*) and fibril density were separated.
- Adhesion areas of cells decreased with increasing fibril density at constant *C*-*E*.
- Adhesion areas of cells increased with increasing *C*-*E* in absence of Col-I fibrils.
- Adhesion areas remained unchanged when both *C*-*E* and fibril density increased.
- Structural, mechanical and topographical cues work in combination to affect cells.

**Graphical Abstract:** 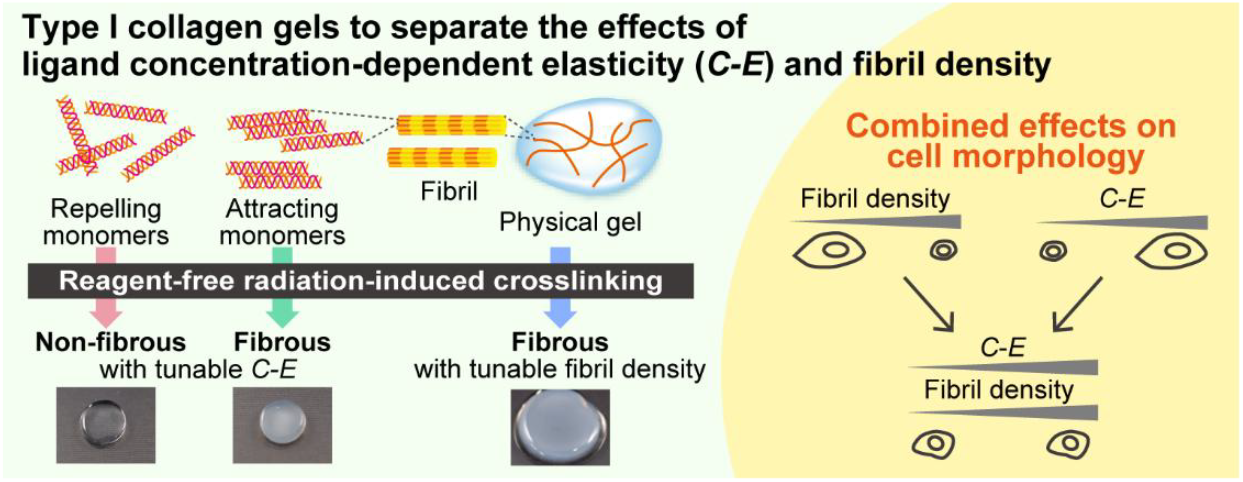

## 1. Introduction

Type I collagen (Col-I) is the most abundant component of the extracellular matrix (ECM), a hydrogel of proteins and polysaccharides. Col-I has hierarchical structures from nanoscale to microscale; its triple helical monomers (tropocollagen, diameter ∼1.5 nm, length ∼300 nm) become fibrils (diameter ∼100 nm, length up to micrometer range) via a self-assembly process called fibrillogenesis, followed by assembling of the fibrils into fibers (diameter ∼10 *μ*m, length up to the millimeter range) via enzyme-mediated crosslinking [1,2]. The mechanical properties of ECM are attributed to these organized architectures of Col-I, as a power law dependence has been reported between the Col-I concentration (from 10 to 200 mg/mL [3]) and the elastic modulus of the bulk soft tissue (from 1 to several hundred kPa) [4,5]. Although the organ-specific composition of ECM is determined during the early embryonic stages, the resident cells produce ECM components and ECM-modifying enzymes constantly, remodeling the biochemical and mechanical properties of the surrounding ECM [6]. For example, in the progression of cancer, fibrosis, vascular diseases, and chronic diseases, as well as aging, it is known that ECM is abnormally stiffened due to the densification of fibrous collagens, mainly Col-I [7]. As the compositional, mechanical, and topographical cues are known to influence the cell functions and fate such as cell migration, proliferation, and differentiation [8–10], the cells surrounded by the altered ECM are affected by the Col-I densification and the accompanying changes in the ECM elasticity and fibril density. Elucidating the effects of these microenvironmental changes on cells will contribute to understanding the mechanisms of various diseases and aging, and to this end, artificial Col-I hydrogels with tunable bioactive ligand concentration, elasticity, and fibril density are required.

ECM-mimicking Col-I gels can be obtained *in vitro* by reconstituting fibrils from triple helical monomers extracted from ECM, under physiological conditions (pH, ionic strength, and temperature) [2]. However, these physical gels self-assemble from low-concentration monomer solutions (< 10 mg/mL) and are fragile, with a typical Col-I concentration (*C*) being approximately 1–2% (w/w) and a resultant gel elasticity (*E*) of only a few kPa. Controlling the *C* and *E* of Col-I gels requires crosslinking between the Col-I molecules, however, conventional techniques require crosslinking agents such as aldehydes or carbodiimides, which consume arginine (R), glutamic acid (E), and aspartic acid (D) residues, thereby damaging the cell binding GxOGER (where G is glycine, O is hydroxyproline, and x is hydrophobic amino acid) motifs in Col-I [11–13]. Another limitation of Col-I physical gels is that gelation is accompanied by fibril formation, therefore, the compositional, mechanical, and topographic cues attributed to fibrils cannot be evaluated separately. To separate the effects of stiffness and fibril density, a pioneering study has developed composite gel of Col-I and non-fibrous gelatin methacrylate to investigate the cellular responses [14]. They demonstrated the importance of evaluating the individual effects of the microenvironmental features on cell behavior, however, it should be noted that Col-I and its hydrolysate, gelatin, have different cell-binding motifs due to structural differences (Col-I: GxOGER, gelatin: RGD) [15]. To overcome this concern, Col-I gels are required, with widely tunable fibril densities while maintaining the triple helical structure, i.e., the cell-binding motifs specific to Col-I.

In this study, to address these challenges, we designed three types of Col-I gels, “A-γ”, “N-γ”, and “P-γ”, that allow separate investigation of the effects of ligand concentration (*C*)-dependent elasticity (*E*) (*C-E*) and fibril density. These gels were produced by introducing crosslinking while adjusting the distance between the Col-I monomers or fibrils, as well as employing the reagent-free radiation-induced crosslinking technique using γ-rays. Both A-γ and N-γ are Col-I gels with controllable *C*-*E*. While N-γ is fibrous, which is the natural characteristic of Col-I, the formation of fibrils in A-γ is difficult. A-γ and N-γ were prepared by crosslinking electrostatically repulsive or attractive monomers in aqueous solutions, respectively. Conversely, P-γ is a fibrous Col-I gel with a tunable fibril density and stable *C*-*E*. P-γ was obtained by crosslinking fibrils in a Col-I physical gel (P). Notably, these gels retained their triple helical structure, i.e., the cell adhesion sequence specific to Col-I. Furthermore, HeLa human cervical cancer epithelial cells, human induced pluripotent stem (hiPS) cells, and 3T3-Swiss albino mouse embryonic fibroblasts were cultured on these Col-I gels and the morphological changes in response to the *C*-*E* and fibril density were investigated. The developed Col-I gels enabled the assessment of the combined effects of compositional, mechanical, and topographical cues of Col-I on cells.

## 2. Materials and Methods

### 2.1. Zeta potential measurement

Acid-soluble Col-I derived from porcine tendon (Cellmatrix Type I-A, 3 mg/mL, pH 3) was obtained from Nitta Gelatin. The zeta potential of acidic and neutralized Col-I solutions was measured using a zeta potential analyzer (ELSZneo; Otsuka Electronics Co., Ltd.) at 10 °C. A Col-I acidic solution (1 mg/mL, pH 3) was prepared by diluting the original 3 mg/mL solution with deionized water (supplied by a Millipore Milli-Q system). The neutralized Col-I solution (pH 7.4) with a physiological ionic strength was prepared by mixing the 1 mg/mL Col-I solution with 10× concentrated phosphate-buffered saline (PBS) and 0.02 M NaOH at a volume ratio of 8:1:1 on ice. The zeta potential was calculated as the average of six measurements (5 runs for each measurement).

### 2.2. Sample preparation

The P-γ, N-γ, and A-γ gels as well as the conventional physical Col-I gel, P, were formed in 35-mm cell culture dishes (3000-035; AGC Techno Glass) or 12-well plates (3815-012; AGC Techno Glass), unless otherwise stated. γ-ray irradiation was conducted at the ^60^Co No. 2 Irradiation Facility of Takasaki Institute for Advanced Quantum Science, QST. The irradiation dose is mentioned after the sample name in kGy (= J/g); for example, 15 kGy-irradiated P-γ samples are indicated as P-γ15.

The physical gel, P, was prepared by mixing the acidic Col-I solution (3 mg/mL, pH 3) with 10× PBS and 0.02 M NaOH at a volume ratio of 8:1:1 on ice. This neutralized solution (pH 7.4) with a physiological ionic strength was dispensed in cell culture dishes or microplates, and heated at 37 °C for 90 min to allow physical gelation of Col-I. P-γ samples were obtained by irradiating P with γ-rays in a styrofoam box containing a heat storage material (sample temperature: ∼30 °C) and subsequently heating at 37 °C for 90 min. A-γ and N-γ were prepared by γ-ray irradiation of the acidic and neutralized Col-I solutions, respectively. These solutions were dispensed in dishes or microplates and were irradiated in a styrofoam box containing ice packs (sample temperature: ∼4 °C). The irradiated A-γ samples were neutralized by adding 10× PBS and NaOH as described above and were subsequently heated at 37 °C for 90 min. The irradiated N-γ samples were directly heated at 37 °C for 90 min. For all Col-I gels, the sample surfaces were covered with polyethylene terephthalate films prior to irradiation, and the films were removed before using the gels.

Crosslinked gelatin gels (Gel-γ) were also prepared and compared with the Col-I gels. Acid-treated gelatin from porcine skin (639-37745, Nitta Gelatin) was dissolved in deionized water at 10 wt% by heating at 50 °C for 30 min. After pouring the gelatin solution into 35-mm cell culture dishes, the samples were sealed in plastic bags with an oxygen absorber (A-500HS, AS ONE) and stored overnight at 20 °C to undergo physical gelation. The gelatin physical gels were irradiated with γ-rays at 15–20 °C for crosslinking. The crosslinked gelatin gels were incubated in PBS at 37 °C for 2 h to remove any non-crosslinked components and allow them to reach equilibrium.

### 2.3. Characterization of the obtained gels

The fibril formation kinetics of Col-I were evaluated from the turbidity of each gel. The gels (final volume of 125 *μ*L) were prepared in each well of round-bottom 96-well plates (449824; Thermo Fisher Scientific), and the optical density (*OD*) was recorded at 405 nm, 37 °C, and 1 min intervals for 90 min using a microplate reader (Multiskan™ FC with incubator, 51119150; Thermo Fisher Scientific). The average blank control (*OD* of the blank wells) was subtracted from the obtained values for each sample.

The microstructures of the Col-I gels were imaged via scanning electron microscopy (SEM) and second-harmonic generation (SHG) microscopy. For SEM, the gels were fixed with 4% paraformaldehyde (163-20145; FUJIFILM Wako Chemicals) overnight. After washing with deionized water, the gels were dehydrated with a series of 25, 50, 70, 90, and 100% ethanol solutions for more than 15 min in each solution. Then, the gels were transferred to 100% hexamethyldisilazane (HMDS) and subsequently dried overnight at room temperature (∼20 °C). The dried samples were imaged using SEM (SU3500, Hitachi High-Tech Science) at 10 kV and magnification of 10,000× after coating the sample surface with platinum using a sputter coater (JFC-1600, JEOL). For SHG microscopy, the gels were fixed in 4% paraformaldehyde for 15 min at room temperature. After washing with PBS, the SHG images of the gels (50 *μ*m under the surface) were obtained using a multiphoton microscope (A1R MP+; Nikon) with a 25× objective lens (CFI75 Apochromat 25XC W; Nikon) at an excitation wavelength of 950 nm. The intensities of the SHG images were measured by ImageJ, while CT-FIRE [16], an open-source software developed and provided by the University of Wisconsin-Madison, was employed to identify the fibrous structures and analyze their width.

The chemical structures of the Col-I gels were evaluated via Fourier transform infrared (FT-IR) spectroscopy after the samples were vacuum-dried overnight at 30 °C. FT-IR spectra were collected using an IRAffinity-1 S instrument (Shimadzu) with a DuraSampl IR-II single-reflection diamond attenuated total reflection attachment (Smiths Detection) at a resolution of 2 cm^−1^ averaged over 64 scans.

The enzyme-mediated degradation properties of the Col-I gels were analyzed using collagenase (034-22363; Fujifilm Wako Pure Chemical). The surface area of the sample affects the rate of degradation, therefore, the gels were adjusted to a similar size (diameter: ∼10 mm, thickness: ∼1.5 mm). After recording their weight (*W*_0_), the samples were incubated in a 0.05% collagenase solution in PBS at 37 °C. At different time points, the samples were filtered through a stainless steel 200-mesh filter (mesh size: 75 *μ*m) and washed with deionized water, followed by weighing (*W*_t_) and calculation of the percentage of remaining weight (%) as (*W*_t_/*W*_0_) × 100.

The Col-I concentration (*C*) was calculated as (*W*_d_/*W*_s_) × 100, where *W*_s_ is the weight of the swollen gel filtered through a stainless steel 200-mesh filter, and *W*_d_ is the weight of the dried gel after vacuum-drying overnight at 30 °C. The elastic modulus (*E*) of the Col-I and gelatin gels was evaluated using an indentation tester (RE2-33005C; Yamaden). The samples were compressed with a 0.2-N load cell using a plunger (*φ*3 mm) at 50 *μ*m/s. The compressive elastic modulus was then determined from the slope of the obtained stress-strain curves over the liner elastic region.

### 2.4. Cell culture

HeLa human cervical carcinoma epithelial cells (RCB0007, RIKEN BRC Cell Bank) and 3T3-Swiss albino mouse embryonic fibroblasts (RCB1642; RIKEN BRC Cell Bank) were cultured in DMEM (08488-55, Nacalai tesque) supplemented with 10% fetal bovine serum (FBS) (12483020; Gibco), 2 mM L-glutamine (G7513; Sigma-Aldrich), 100 units/mL penicillin, and 100 *μ*g/mL streptomycin (15144122; Gibco). The Col-I or gelatin gels in 35-mm dishes were pre-incubated with 2 mL PBS and 5% CO_2_ at 37 °C for 2 h. After removing PBS, 2 mL of the culture medium was added and pre-incubated at 37 °C in 5% CO_2_ for > 30 min. After removing the medium, 2 mL of the culture medium was added and pre-incubated at 37 °C in 5% CO_2_ for > 30 min. A 2-mL portion of each cell suspension (1×10^4^ cells/mL) was seeded on the Col-I or gelatin gels and incubated at 37 °C in 5% CO_2_ for 3 d.

The hiPS cells (1383D2; HPS1005, RIKEN BRC Cell Bank) were cultured in StemFit AK02N (Ajinomoto Healthy Supply). The gels in 35-mm dishes were pre-incubated with PBS and 5% CO_2_ at 37 °C for 2 h as described above. After removing PBS, 2 mL of StemFit AK02N with 10 *μ*M Y27632 (036-24023; FUJIFILM Wako Chemicals) was added and pre-incubated at 37 °C in 5% CO_2_ for > 30 min twice as described above. The 10^5^ cells in 2 mL of StemFit AK02N with Y27632 were seeded on the Col-I or gelatin gels and incubated at 37 °C in 5% CO_2_ for 1 d.

### 2.5. Fluorescence staining and imaging of cells

The cells on the Col-I or gelatin gels were washed with PBS and fixed in 4% paraformaldehyde in PBS for 15 min at room temperature. After washing with PBS, the cells were incubated for 5 min in PBS containing 0.1% Triton X-100 (35501-02; Nacalai Tesque). After washing with PBS, the cells were incubated in PBS containing 1% bovine serum albumin (BSA) (P-6154; Biowest) and 0.1 *μ*g/mL tetramethylrhodamine B isothiocyanate (TRITC)-conjugated phalloidin (P1951; Sigma-Aldrich) for 20 min at room temperature. The fluorescence images of the stained cells were obtained using a multiphoton microscope with a 25× objective lens at an excitation wavelength of 950 nm. The shapes of the cells were manually traced, and the areas were measured by ImageJ.

### 2.6. Statistical analysis

The statistical significance between the two groups was assessed using Welch’s t-test. For multiple tests, the statistical significance was assessed by one-way ANOVA with Tukey’s post-test. The values of *P* < 0.05 were considered statistically significant. The analysis was performed using OriginPro 2023 (OriginLab). The data are expressed as the mean ± standard error of the mean (s.e.m.) unless otherwise stated.

## 3. Results and Discussion

### 3.1. Design and formation of Col-I gels

Col-I gels were developed by introducing radiation-induced crosslinking while adjusting the distance between the Col-I monomers or fibrils (Fig. 1), to separately evaluate the effects of *C*-*E* and fibril density on cell behavior.

**Fig. 1.**
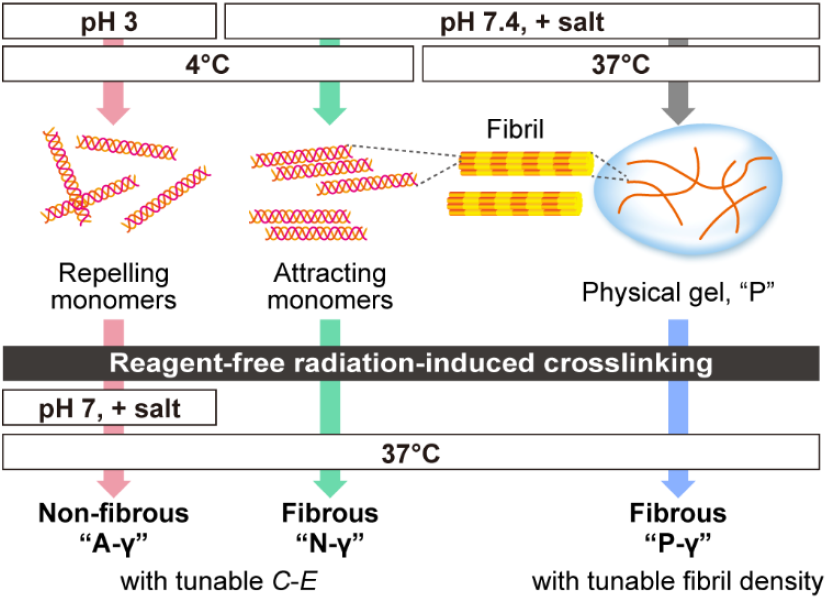
Design, preparation procedure, and targeted properties of the three types of Col-I gels, “A-γ”, “N-γ”, and “P-γ”, enabling the evaluation of the combined effects of *C*-*E* and fibril density on cells.

The aggregation of the Col-I triple helical monomers is driven by pH and ionic conditions that tune the electrostatic interactions, as well as temperature that affects hydrophobic interactions [2]. The zeta potential of the Col-I monomers was ∼26.8 mV in an acidic solution (pH 3) at 10 °C (Table S1), indicating that the monomers electrostatically repelled each other. When PBS and NaOH were added to the acidic Col-I solution to afford a neutral solution (pH 7.4) with a physiological ionic strength, the zeta potential value decreased to nearly zero, ∼0.03 mV (Table S1). Such a finding indicates that the isoelectric point of Col-I in this system is around pH 7.4. The electrostatic repulsion between monomers is minimized in this neutral solution; therefore, upon heating to 37 °C, the monomers self-assembled into fibrils to form a physical gel, P.

Introduction of crosslinks was attempted under three different conditions depending on the distance between the Col-I monomers or fibrils: in an acidic solution where monomers repel each other, in a neutral solution where monomer aggregation is promoted, and in a physical gel after fibril formation. A reagent-free radiation-induced crosslinking method was employed; although Col-I is dominantly decomposed by ionizing radiation in a dried state [17], crosslinking is induced in the presence of water through a reaction with generated hydroxyl radicals [18–21]. Therefore, the Col-I samples were irradiated with γ-rays in an acidic solution, neutral solution, and physical gel, referred to as A-γ, N-γ, and P-γ, respectively (Fig. 1).

The appearance of P-γ did not change significantly compared to that before irradiation (P). Conversely, the acidic and neutral solutions directly changed into hydrogels upon irradiation, and A-γ and N-γ were obtained as transparent and cloudy gels, respectively. The gelation effect of ionizing radiations on the Col-I acidic solution has been reported in several studies [19,21–23]. However, to the best of our knowledge, our study is the first to directly gel a Col-I solution with physiological pH and ionic strength via ionizing radiation. Our previous study on gelatin (hydrolysate of Col-I) showed that OH radicals from water radiolysis introduce stable radicals into the aromatic rings of phenylalanine, tyrosine, and histidine residues, resulting in crosslinking via bimolecular reactions [21,24]. Therefore, we believe that γ-ray-irradiation induced chemical crosslinking of the Col-I molecules in both the acidic and neutral solutions via the same mechanism as gelatin, without requiring reagents or heating.

### 3.2. Changes in Col-I gels under a physiological environment

All the γ-ray-irradiated Col-I gels were further altered under the physiological environment. Fig. 2A shows the samples prepared and irradiated in 12-well plates (diameter: 22 mm) and subsequently heated at 37 °C for 90 min. A-γ samples were adjusted to a similar physiological pH and ionic strength to those of the other two types of gels by adding PBS and NaOH before heating. Compared with the physical gel (P) formed at the 12-well size, A-γ and N-γ shrank significantly with increasing irradiation dose. A-γ was almost transparent even after contraction, however, N-γ exhibited increased turbidity depending on the irradiation dose. Conversely, P-γ did not exhibit any change in its size, although the turbidity increased in accordance with the irradiation dose.

**Fig. 2.**
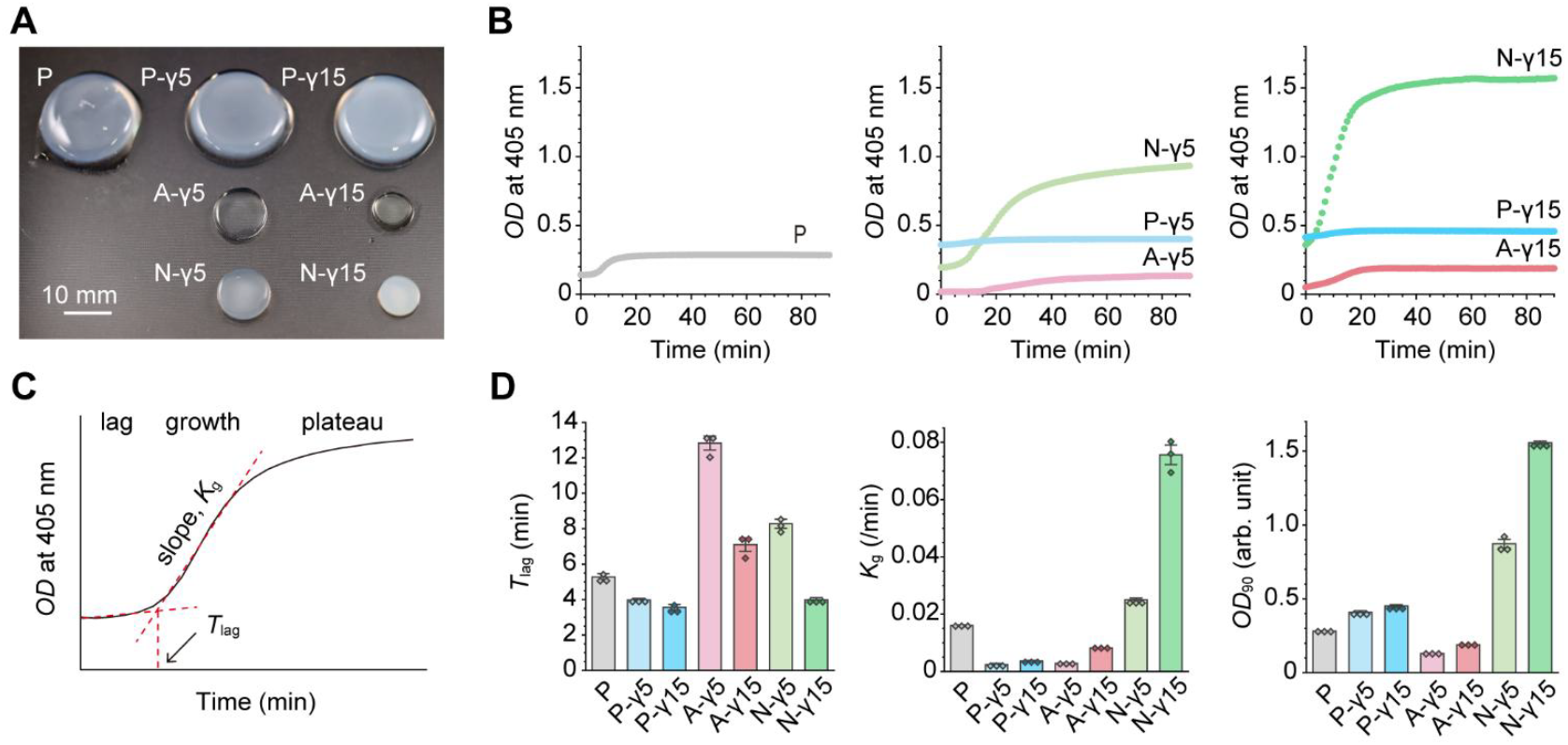
Temporal changes of the Col-I gels under the physiological environment evaluated from the viewpoint of turbidity. (A) A photograph of the P, P-γ, N-γ, and A-γ gels after heated at 37 °C for 90 min. The number after the sample name implies the irradiation dose in kGy. (B) Temporal changes in the turbidity of each gel assessed by monitoring *OD* at a wavelength of 405 nm at 1-min intervals. (C) A typical sigmoid curve drawn by the change of turbidity over time through fibril formation. (D) The time required for switching from the lag phase to the growth phase (*T*_lag_), the slope of the growth phase (*K*_g_), and the *OD* values of the plateau phase after 90 min of heating (*OD*_90_) for each gel (*n* =3).

The temporal changes of the Col-I gels under the physiological environment were evaluated from the viewpoint of turbidity. The measurement of turbidity is a simple and powerful technique for assessing fibrillogenesis [2]. In this study, *OD* was measured at 405 nm and 1-min intervals using a microplate reader heated to 37 °C. As shown in Fig. 2B, the *OD* value of each Col-I gel increased, following a sigmoidal curve over several tens of minutes. Regarding the physical gel (P), it is known that the transition from the lag phase, where the change in turbidity is little, to the growth phase, where the light scattering increases rapidly, and then to the plateau phase, where turbidity is almost stable, corresponds to the process in which the Col-I monomers aggregate and assemble into fibrils [2]. After γ-ray irradiation of the physical gel (P-γ), the turbidity increased at 37 °C, and the degree of change increased depending on the irradiation dose. Such results indicate that γ-ray irradiation further aggregated the fibrils in P. The turbidity of A-γ also increased slightly according to the irradiation dose, however, it was lower than that of P, indicating that fibril formation may be inhibited by the crosslinked structure. Conversely, the turbidity of the N-γ gel exceeded that of P and increased according to the irradiation dose, suggesting that fibrillogenesis or further fibril aggregation was promoted by crosslinking.

The time to switch from the lag phase to the growth phase in which *OD* increases linearly (*T*_lag_), the slope of the growth phase (*K*_g_), and the *OD* values of the plateau phase after 90 min of heating (*OD*_90_) were extracted from the sigmoid curves obtained for each gel, as shown in Fig. 2C. As summarized in Fig. 2D, P-γ was the fastest in entering the growth phase at the same irradiation dose. Conversely, N-γ exhibited the largest growth rate and *OD*_90_ value. In all gels (P-γ, N-γ, and A-γ), the increased irradiation dose shortened the time required to reach the growth phase, accelerated the growth rate, and increased the final turbidity. It was therefore suggested that the increase in the irradiation dose increased the number of crosslinking points, which promoted the structural changes in each gel.

### 3.3. Evaluation of fibril density in Col-I gels

The microstructure of each Col-I gel was observed via SEM (Fig. S1) revealing Col-I fibril structures in the P, P-γ, and N-γ gels, but not in the A-γ gels, as expected from their nearly transparent nature (Fig. 2A). It was assumed that the crosslinks formed between the repulsive monomers inhibited fibril formation in A-γ. In the SEM images, the fibrils appear to be thicker in the order P < P-γ < N-γ; however, it is difficult to evaluate the fibril shape since the aggregation and shrinkage of fibrils are inevitable during the drying process.

Therefore, we next employed SHG microscopy, which allows observation of swollen gels. Similarly to SEM (Fig. S1), fibrous structures were observed for P, P-γ, and N-γ (Fig. 3A). A correlation was observed between the SHG and SEM images, however, the observed fibril diameter was different by a factor of ∼10. Such a difference was attributed to two factors: the thickness of the fibrils in the SEM images appears thinner due to the drying process, furthermore, the SHG images do not necessarily show the exact fibril shape because the intensity of the SHG images depends on the shape and size of the structures [25]. Therefore, the SHG images were used to evaluate the space-filling properties of the fibril structure of the Col-I gels, i.e., the fibril density. The SHG intensities measured by ImageJ and width of the fibrillar structures automatically extracted from the SHG images (*W*_SHG_) using CT-FIRE [16] both increased with increasing irradiation dose (Fig. 3B, C). Such results indicate that the fibril density in the P-γ and N-γ gels increased via crosslinking. It was assumed that in P-γ, the formation of crosslinks between fibrils led to fibril aggregation, whereas in N-γ, the crosslinks between attracting monomers promoted fibril formation and aggregation.

**Fig. 3.**
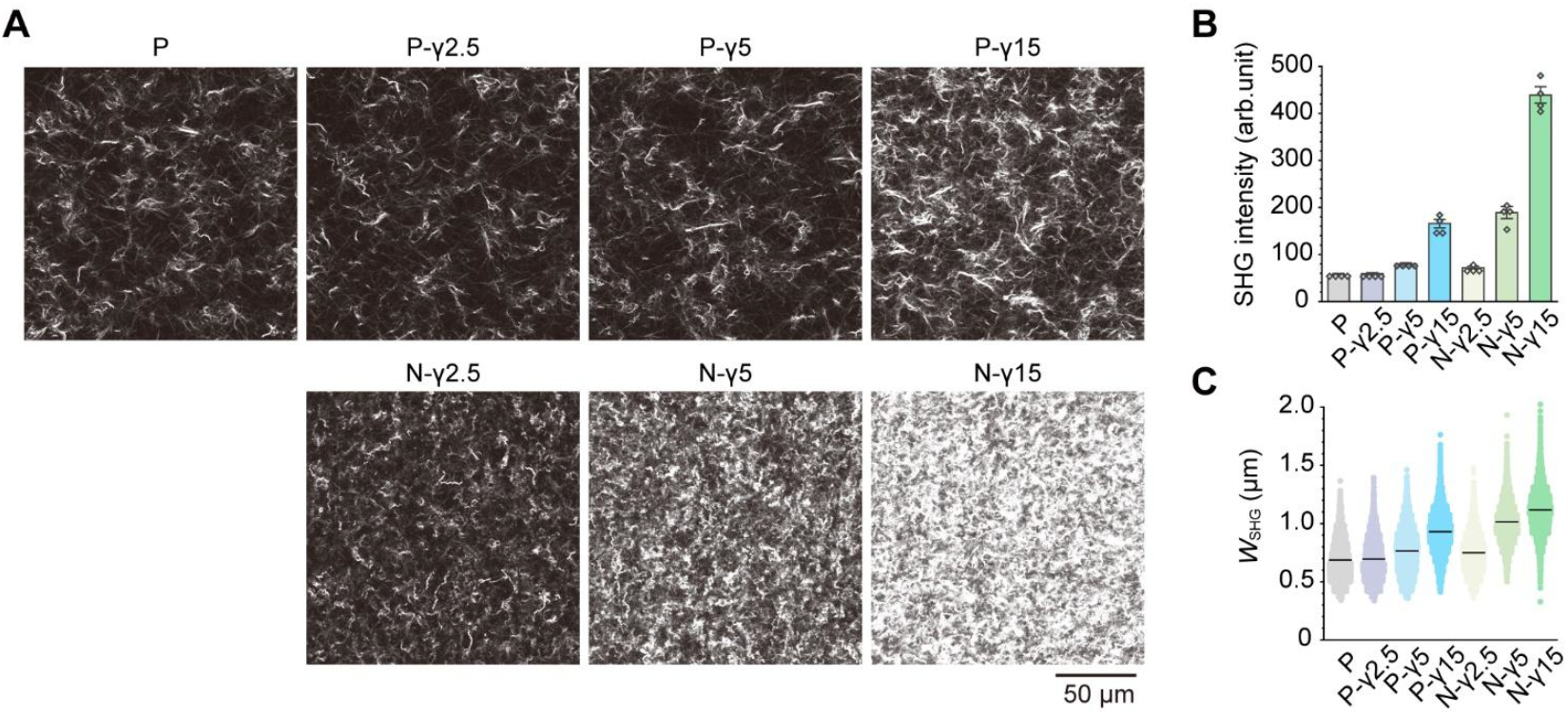
Evaluation of the fibril density of Col-I gels. (A) Representative SHG images of each gel. (B) Intensity of the SHG images. (C) Width of the structures automatically extracted by CT-FIRE from the SHG images (*W*_SHG_). In (B) and (C), four SHG images were analyzed for each condition. The bars in (C) indicate the mean values; the numbers of extracted structures for P, P-γ2.5, P-γ5, P-γ15, N-γ2.5, N-γ5, and N-γ15 were 3752, 3817, 4094, 5018, 4705, 5020, and 4686, respectively.

### 3.4. Chemical and mechanical characteristics of Col-I gels

The dried Col-I gels were analyzed via FT-IR to investigate the effect of crosslinking on the Col-I molecules (Fig. 4A). All the samples showed characteristic absorption bands related to the peptide linkages of Col-I: amide A at 3300 cm^−1^ and amide B at 3070 cm^−1^ mainly associated with the N–H stretching vibrations; amide I at 1640 cm^−1^ corresponding to the C=O stretching vibrations; amide II at 1536 cm^−1^ consisting of the N–H bending and C–N stretching vibrations; and amide III at 1236 cm^−1^ assigned to the C–N stretching and N–H bending vibrations [26]. No clear difference was observed between the spectra obtained for each gel, indicating that the peptide linkages of the Col-I molecules were essentially unaltered by the introduction of crosslinking and the formation of gels. Such a result is consistent with our previous findings on gelatin being crosslinked and gelled by a radiation-induced reaction; γ-ray irradiation up to 60 kGy did not afford significant changes in the amino acid composition of gelatin [21,24]. Conventional methods use aldehydes and carbodiimide, as well as consume R, E, and D, thus damaging cell adhesion motifs in both gelatin (RDG) and collagen (GxOGER) [11–13]. Compared with such methods, our crosslinking approach is superior, preserving the amino acid composition.

**Fig. 4.**
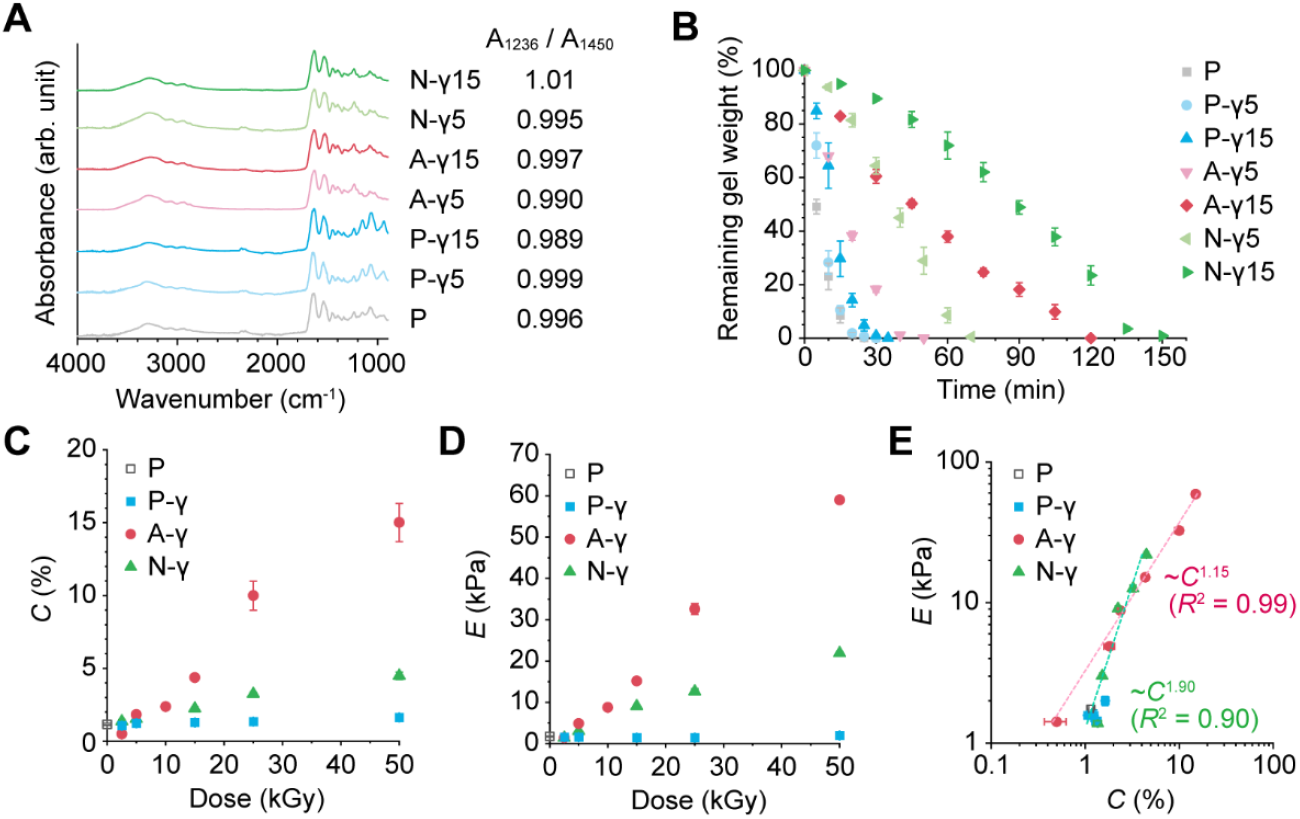
Chemical and mechanical characterization of the Col-I gels. (A) Representative FT-IR spectra of the obtained Col-I gels and the absorption ratio of Amide III (1236 cm^−1^) to 1450 cm^−1^ (*A*_1236_/*A*_1450_), which is an indicator of the conservation of the triple helix structure of Col-I. (B) Enzymatic degradation test using collagenase (*n* = 4). (C) Concentration of the Col-I constitutes in the gels, *C* (*n* = 3) (D) Elastic moduli of the Col-I gels, *E*. Three samples were measured at three points each. (E) Dependence of *E* on *C*.

To analyze the preservation degree of the triple helical structure of Col-I, the absorption ratio of amide III (1236 cm^−1^) to 1450 cm^−1^, *A*_1236_/*A*_1450_, was calculated. According to previous reports, this ratio is close to 1 when the triple helical structure is preserved, and decreases to ∼0.8 when the triple helix is denatured (unfolded or misfolded) [26]. According to Fig. 4A, the *A*_1236_/*A*_1450_ values for all the Col-I gels were almost 1, indicating that the triple helical structure of the Col-I molecule was largely preserved in all gels (P-γ, A-γ, and N-γ). Therefore, the major cell adhesion sequence in all Col-I gels is considered to be GxOGER, which is derived from the triple helical structure.

The enzymatic degradability of the obtained gels was evaluated using collagenase. As shown in Fig. 4B, the Col-I gels retained their enzymatic degradability. This result is consistent with the FT-IR analysis (Fig. 4A), indicating that the chemical structure of Col-I remains unchanged. Gels of the same type exhibited decreased degradation rates with increasing irradiation dose, that is, crosslinking density. Conversely, gels of different types exhibited degradation rates in the order P > P-γ > A-γ > N-γ, and tended to decrease with increasing fibril density (Fig. 3) or the concentration of Col-I, *C* (Fig. 4C).

*C* and the compressive elastic modulus of the gels, *E*, are summarized in Fig. 4C, D. *C* and *E* of A-γ and N-γ increased with increasing irradiation dose, i.e., increasing the number of crosslinking points, however, *C* and *E* of P-γ hardly changed and were comparable to those of P. This result is consistent with the conclusion that the A-γ and N-γ gels significantly shrank, while the size of the P-γ gels did not change (Fig. 2A). Regarding P-γ, since fibrils are already formed via physical gelation, it is thought that bulk dehydration and condensation, and the associated increases in *C* and *E*, are unlikely to occur. A power-law correlation was observed between *C* and *E* of A-γ and N-γ (Fig. 4E), which is similar to that reported between the *C* of Col-I in native ECM and *E* of bulk soft tissues [5].

Such findings confirmed the formation of three Col-I gels with different features. P-γ is a fibrous Col-I gel in which the fibril density can be tuned without changing *C*-*E*, making it suitable for investigating the effect of fibrils separately from other cues. Both A-γ and N-γ are Col-I gels with tunable *C*-*E*. In addition to the effects of *C*-*E* from non-fibrous A-γ and fibrous N-γ, the influence of the fibril density can be investigated by comparing the cellular response to these two types of gels. N-γ, in which *C*-*E* and the fibril density change simultaneously, is thought to best reproduce the changes in the microenvironment that accompany the densification of Col-I *in vivo*. Importantly, P-γ, A-γ, and N-γ retained the triple helical structure of Col-I, i.e., the cell-binding motifs specific to Col-I.

### 3.5. Cell culture on Col-I gels

The developed three types of Col-I gels allowed us to investigate the individual and combined effects of *C*-*E* and fibril density on cells. HeLa cervical cancer epithelial cells, hiPSCs, and 3T3-Swiss fibroblasts were therefore cultured on the developed Col-I gels to analyze the cell adhesion area.

First, the effect of fibril density was investigated alone. Fig. 5A shows the fluorescence images of actin filaments in cells cultured on P and P-γ with different fibril densities and constant *C*-*E* (*C* = ∼1.2%, *E* = ∼1.5 kPa). The adhesion area of the cells is summarized in Fig. 5B. All cells changed their morphology depending on the fibril density. HeLa cells spread and formed a well-developed actin cytoskeleton on P and P-γ2.5, however, the adhesion area became smaller as the fibril density increased. hiPS cells showed a similar but more drastic change and only spread well on P. Regarding 3T3-Swiss cells, they exhibited the largest adhesion area on P-γ2.5, while in general, their adhesion area decreased as the fibril density increased. The change in adhesion area due to the change in fibril density was smaller than that of the other two cell types.

**Fig. 5.**
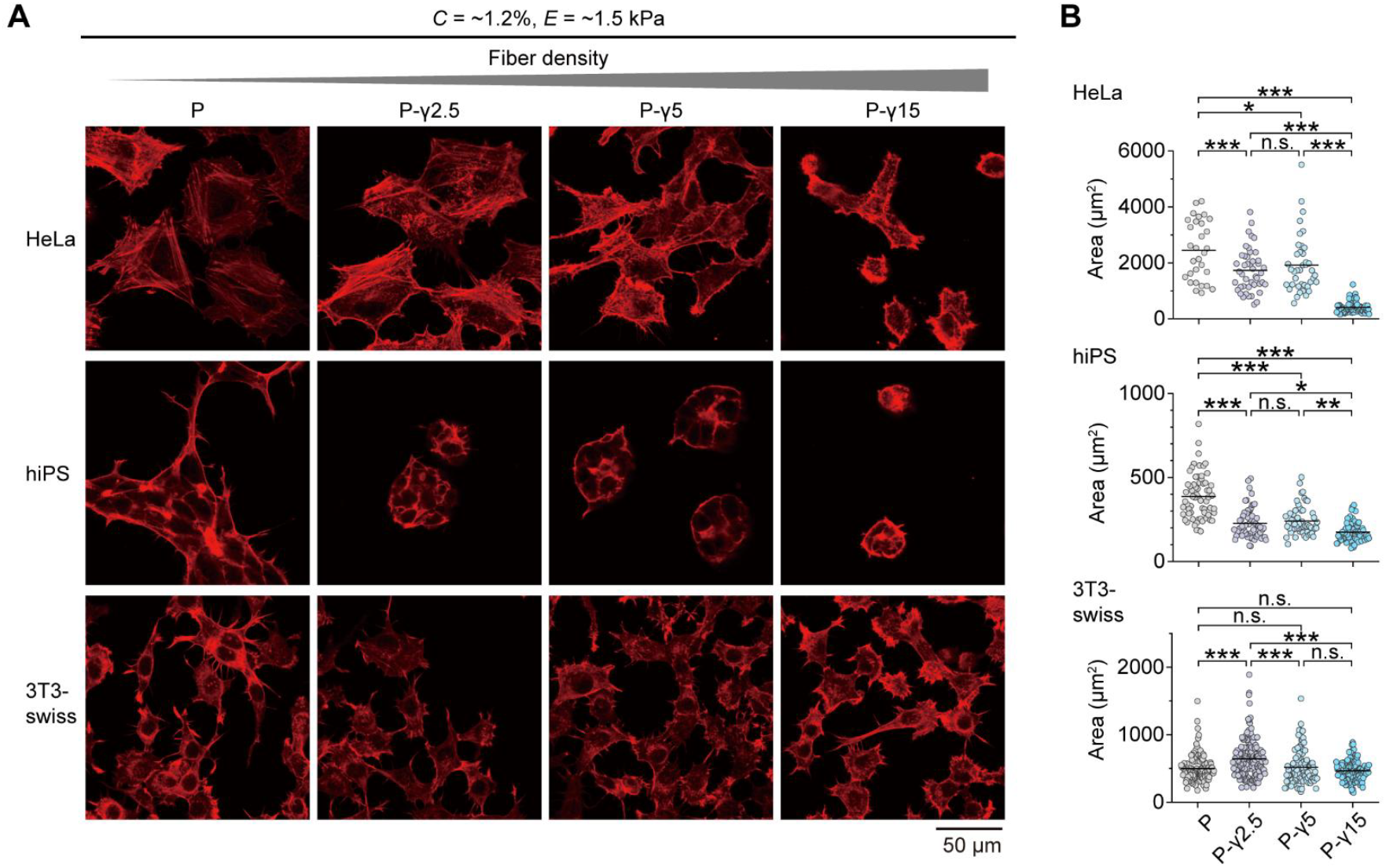
Effect of the fibril density of the Col-I gels on cell morphology. (A) Fluorescence images of actin filaments and (B) cell area of HeLa, hiPS, and 3T3-swiss cells cultured on Col-I gels with constant *C*-*E* (*C* = ∼1.2%, *E* = ∼1.5 kPa) and different fibril densities. The bars in (B) indicate the mean values; the statistical significance was assessed by one-way ANOVA with Tukey’s post-test. n.s., not significant (P ≥ 0.05); *, P < 0.05; **, P < 0.01; ***, P < 0.001. See Table S2 for the *n* and P values.

Next, the cell morphology was compared on A-γ and N-γ gels with tunable *C*-*E*. For non-fibrous A-γ, only the effect of *C*-*E* was considered, whereas for fibrous N-γ, the fibril density increased with *C*-*E*.

A-γ10 and N-γ15, which have similar *C*-*E* (*C* = ∼2.3%, *E* = ∼9.0 kPa), were compared first. As shown in Figs. 6A and 6C, the adhesion area of HeLa and 3T3-swiss cells decreased on N-γ15, which has a higher fibril density compared with the non-fibrous A-γ10. This is consistent with the reduced cell area observed with increasing fibril density on P and P-γ (*C* = ∼1.2%, *E* = ∼1.5 kPa, Fig. 5). Next, A-γ2.5 (*C* = ∼0.5%, *E* = ∼1.4 kPa) was compared with A-γ10 (*C* = ∼2.3%, *E* = ∼9.0 kPa) to investigate the effect of *C*-*E* without the influence of fibrils. The adhesion area of both HeLa and 3T3-swiss cells increased with increasing *C*-*E* (Figs. 6A and 6C), similar to our previous report using non-fibrous crosslinked gelatin gels [21]. Notably, the cell adhesion area on N-γ2.5 and N-γ15, in which *C*-*E* and the fibril density increased simultaneously, did not change for both HeLa and 3T3-swiss cells (Figs. 6B and 6C).

**Fig. 6.**
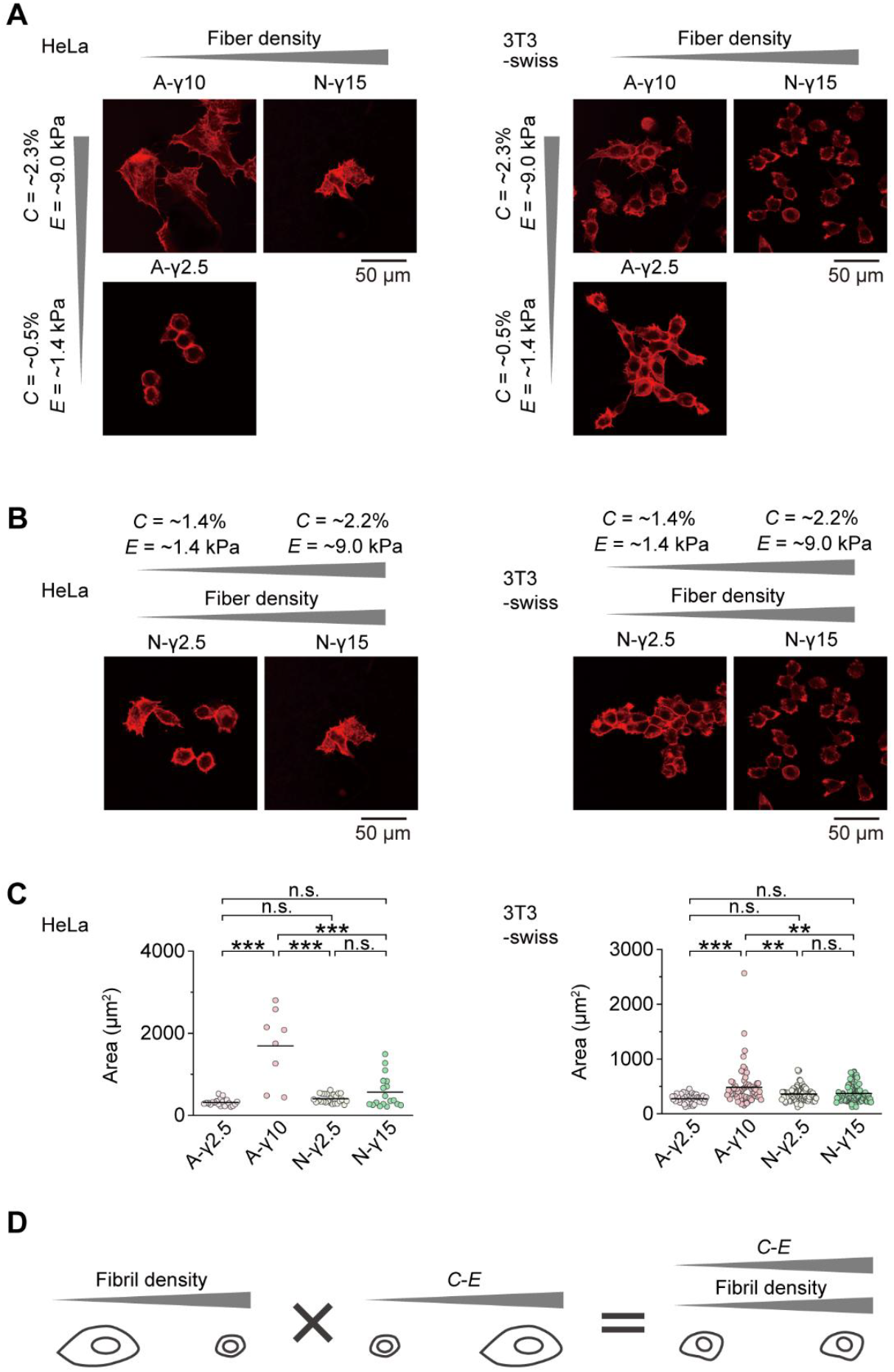
Combined effects of *C*-*E* and the fibril density of Col-I gels on cell morphology. (A) Fluorescence images of actin filaments in HeLa and 3T3-swiss cells cultured on Col-I gels with different *C*-*E* or different fibril densities. (B) Fluorescence images of actin filaments in the cells cultured on Col-I gels with simultaneously varying *C*-*E* and fibril density. (C) Summary of the cell adhesion areas. (D) Representation of the changes in the cell adhesion area in response to the combined effects of *C*-*E* and fibril density of Col-I gels. The bars in (C) indicate the mean values; the statistical significance was assessed by one-way ANOVA with Tukey’s post-test. n.s., not significant (P ≥ 0.05); *, P < 0.05; **, P < 0.01; ***, P < 0.001. See Table S2 for the *n* and P values.

HeLa and 3T3-Swiss cells adhered and spread on all Col-I gels, including the non-fibrous A-γ, allowing us to verify and summarize the combined effects of *C*-*E* and fibril density on the cell adhesion area (Fig. 6D). At a constant *C*-*E*, the cell adhesion area decreased with increasing fibril density (P and P-γ in Fig. 5 and A-γ10 vs N-γ15 in Figs. 6A and 6C). In the absence of the fibrils effect, the cell adhesion area increased with increasing *C*-*E* (A-γ2.5 vs A-γ10 in Figs. 6A and 6C). When *C*-*E* and the fibril density increased simultaneously, mimicking the changes in the *in vivo* ECM caused by Col-I densification, the two opposing effects counteract each other and the cell adhesion area remains unchanged (N-γ2.5 vs N-γ10 in Figs. 6B and 6C). The response of hiPS cells was different from that of HeLa and 3T3-Swiss cells; they formed clusters on the non-fibrous A-γ gels regardless of *C*-*E* (Fig. S2). Thus, it was concluded that hiPS cells adhere two-dimensionally only within a certain range of fibril densities (Fig. 5).

We also prepared the crosslinked gelatin gels (Gel-γ6) with the same *E* as A-γ10. Both A-γ10 and Gel-γ6 are non-fibrous, exhibiting differences in their cell-binding motifs (Col-I: GxOGER, gelatin: RGD) [15]. hiPS cells formed clusters on both gels, however, HeLa and 3T3-swiss cells spread more on A-γ10 gels, sensing the difference in the composition of the contact surface (Fig. S3). Such a result highlights the importance of the developed gels, which are made only from Col-I and can control the ligand concentration, elasticity, and fibril density.

To the best of our knowledge, this is the first study to investigate the combined effects of *C*-*E* and fibril density of Col-I gels on cells, although there have been several reports on the combined effects of *C* and topography, or *E* and topography. For example, it has been reported that *C* and topography have an additive effect on the adhesion of HepG2 cells using a microtextured poly(DL-glycolic-co-lactic acid) copolymer coated with Col-I [27]. Similarly, it has been revealed that the combination of *C* of laminin and submicron to micropillars made of silicon promotes the adhesion of mouse embryonic stem cells [28]. Regarding the combined effect of *E* and topography, several reports have shown that the improvement in cell spreading due to an increase in *E* observed on a flat surface can be reversed by the effect of topography [29–31]. In particular, Baker et al. used fibers made of dextran methacrylate and showed that increasing the *E* of the fiber inhibited the spreading and proliferation of human mesenchymal stem cells [29]. They proposed a mechanism of “fiber recruitment” in which cells recruit fibers with low *E*, which increases *C* on the cell surface and promotes the formation of focal adhesions and associated signaling. Xie et al. prepared Col-I gels with fibrils of different *E* while keeping *C* constant, and showed that cell spreading was inhibited on gels with stiff fibrils [32].

Considering the aforementioned studies, we believe that the combined effect of *E* and topography is dominant in the cancellation of the effects of *C*-*E* and fibril density on the cell adhesion area. In the absence of fibrils, increases in *C*-*E* worked in the direction of cell spreading. In this study, we achieved the development of a non-fibrous Col-I gel, A-γ, by inhibiting fibril formation, allowing us to examine only the effect of *C*-*E* of Col-I. However, when the fibril density increased, cell spreading was inhibited regardless of *C*-*E*. The increased fibril density reduced the number of loose fibrils that cells could recruit, potentially negatively affecting cell spreading. The developed Col-I gels allowed the separate assessment of the effects of *C*-*E* and fiber density, revealing that the simultaneous increase in *C*-*E* and fiber density that would occur upon Col-I densification *in vivo* counteracts the effects on cell spreading.

Although more detailed investigations are required, the obtained results suggest the existence of an optimal fibril density for cell spreading depending on the cell type. HeLa and hiPS cells spread most widely at P, whereas 3T3-swiss cells spread at P-γ2.5. hiPS cells were particularly sensitive to Col-I fibril density, exhibiting clustering without spreading outside a specific fibril density range, regardless of *C*-*E*. Furthermore, the extent of the influence of Col-I *C*-*E* and fibril density varied depending on the cell type. 3T3-swiss cells were less affected by both *C*-*E* and fibril density than the other two types of cells, which may be because they are Col-I producing cells.

## 4. Conclusions

Col-I gels were designed to allow investigation of the combined effects of microenvironmental changes caused by Col-I densification, i.e., increased *C*-*E* and fibril density, on cells. By introducing reagent-free radiation-induced crosslinking between the Col-I monomers or fibrils, we fabricated three types of gels: P-γ, which has a constant *C*-*E* and adjustable fibril density; A-γ, which is non-fibrous and has adjustable *C*-*E*; and N-γ, with tunable *C*-*E* and the fibril density simultaneously. FT-IR analysis revealed that all the developed gels retained the triple helical structure of Col-I, i.e., its original cell-binding motifs. HeLa, hiPS, and 3T3-Swiss cells were cultured on these gels and the cell morphology was observed, revealing that the adhesion area of all cells decreased with increasing fibril density when *C*-*E* was constant. In the absence of fibrils, HeLa and 3T3-Swiss cells showed an increase in adhesion area with increasing *C*-*E*, however, hiPS cells, which only spread on gels with a specific fibril density, formed clusters regardless of *C*-*E*. When *C*-*E* and the fibril density were increased simultaneously, i.e., mimicking the changes in the native ECM caused by Col-I densification, the adhesion area of the cells did not change because of the two opposing effects. Our results highlight the importance of elucidating the combined effects of Col-I compositional, mechanical, and topographical cues on cells. The developed gels can provide insights into the interactions between Col-I and cells, to contribute to early detection of diseases, disease progression testing, treatment, prognosis prediction, tissue engineering, and regenerative medicine.

## Declaration of Competing Interest

T.G.O., K.O., and M.T. are co-inventors on filed patent applications related to this work.

## Acknowledgements

The authors thank Ms. Ryoko Mezaki (QST), Ms. Noriko Tawara (QST), Ms. Yukiko Sakatsume (QST), and Ms. Yoko Shimoyama (QST) for their technical assistance. We would like to thank Editage for English language editing.

## Funding

This study was supported by ACT-X [grant number JPMJAX2014]; A-STEP [grant number JPMJTR22U7] from the Japan Science and Technology Agency (JST); Innovative Science & Technology Initiative for Security [grant number JPJ004596] from Acquisition, Technology & Logistics Agency (ATLA); and KAKENHI [grant number 24K01998] from Japan Society for the Promotion of Science (JSPS).

## Author contributions

T.G.O. conceived and initiated the project. T.G.O. and K.O. designed and performed the experiments and analyzed the data. T.G.O., K.O., H.M., and M.T. wrote the manuscript.

## Data availability

The data that support the findings of this study are available from the corresponding authors upon reasonable request.

## Supplementary material

Supplementary data to this article can be found online.

