## Supplementary material for "Type I collagen gels for assessing the combined effects of ligand concentration-dependent elasticity and fibril density on cells"

Table S1. Zeta potential of the Col-I solutions measured at 10 °C.

| | Zeta potential (mV) | | | | | | | |
|---|---|---|---|---|---|---|---|---|
| | 1 | 2 | 3 | 4 | 5 | 6 | mean | s.e.m. |
| pH 3, without salt | 26.29 | 25.52 | 27.32 | 26.97 | 27.15 | 27.55 | 26.8 | 0.31 |
| pH 7.4, with salt | −1.07 | −0.77 | −0.38 | 0.28 | 1.03 | 1.11 | 0.03 | 0.38 |

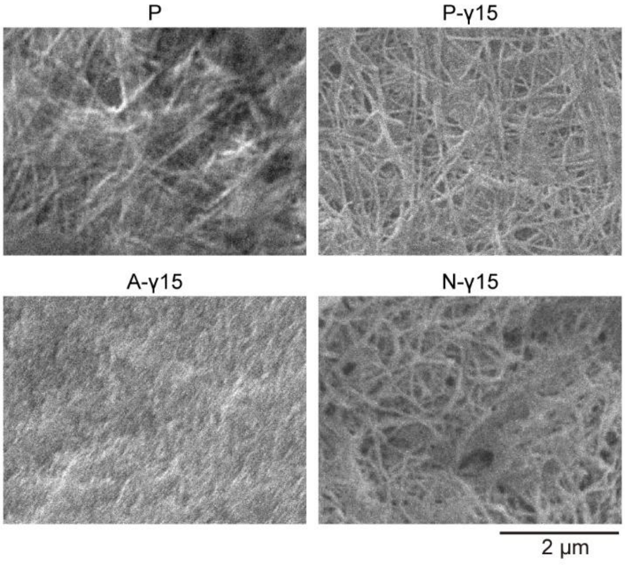


Fig. S1 Representative SEM images of each Col-I gel.

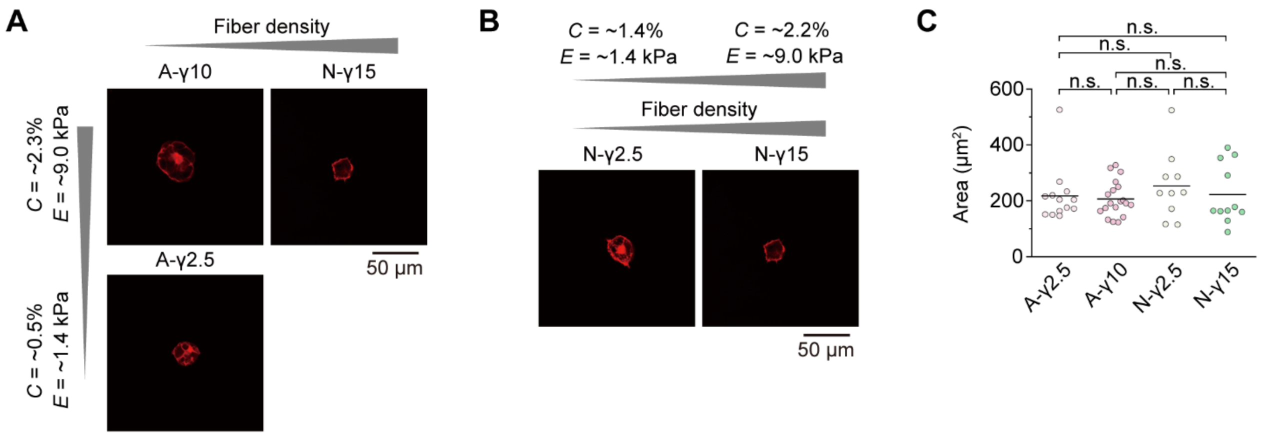


Fig. S2 Individual or combined effects of *C*-*E* and fibril density of Col-I gels on cell morphology. (A) Fluorescence images of actin filaments in hiPS cells cultured on Col-I gels with different *C*-*E* or fibril densities. (B) Fluorescence images of actin filaments in the cells cultured on Col-I gels upon simultaneously varying *C*-*E* and the fibril density. (C) Summary of the cell adhesion areas. The bars in (C) indicate the mean values; the statistical significance was assessed by one-way ANOVA with Tukey's post-test. n.s., not significant ($P \geq 0.05$); *, $P < 0.05$; **, $P < 0.01$; ***, $P < 0.001$. See Table S2 for the *n* and P values.

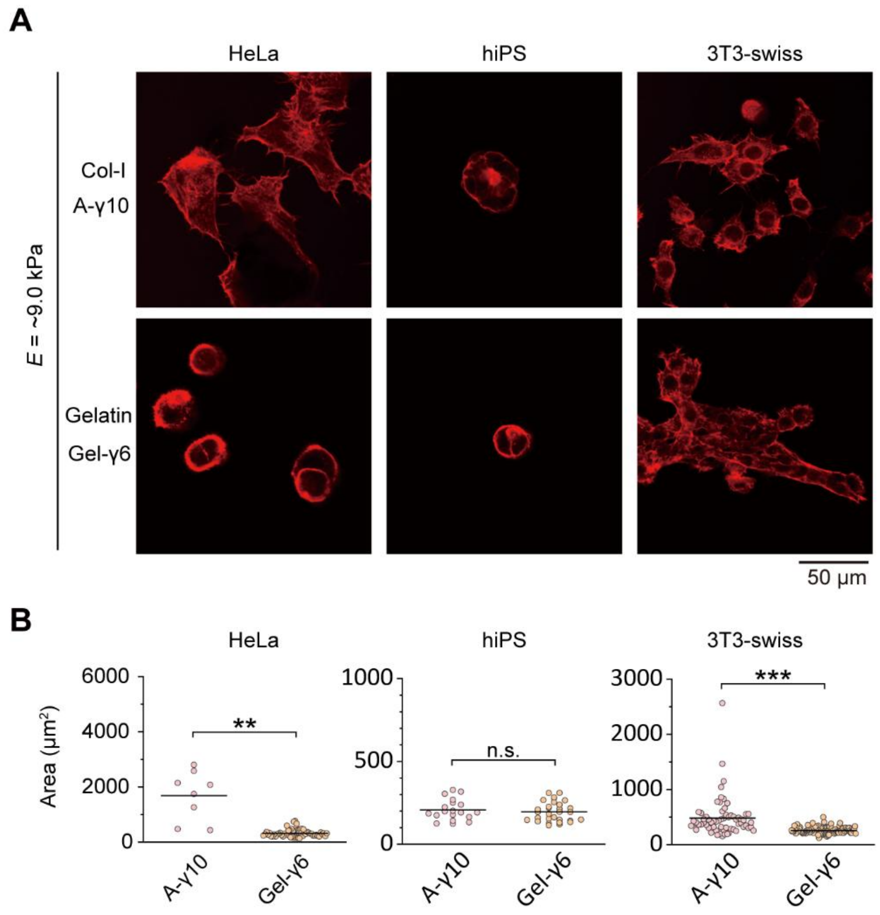


Fig. S3 (A) Fluorescence images of actin filaments and (B) cell areas of HeLa, hiPS, and 3T3-swiss cells cultured on A-γ10 made of Col-I and Gel-γ6 made of gelatin, having the same elasticity (*E* = ~9.0 kPa). The A-γ10 images are the same as those presented in Fig. 6 and Fig. S2, and are shown here for comparison with the Gel-γ6 images. The bars in (B) indicate the mean values. The statistical significance was assessed by Welch's t-test. n.s., not significant ($P \geq 0.05$); *, $P < 0.05$; **, $P < 0.01$; ***, $P < 0.001$. See Table S2 for the *n* and P values.

Table S2. Datasets of the cell responses analyzed in this study.

| Related Figure | Cell type | n | | P | |
|---|---|---|---|---|---|
| Fig. 5B | HeLa | P | 30 | P-γ2.5 vs P | 1.90E-04 |
| | | P-γ2.5 | 45 | P-γ5 vs P | 0.01406 |
| | | P-γ5 | 40 | P-γ5 vs P-γ2.5 | 0.60889 |
| | | P-γ10 | 89 | P-γ15 vs P | < 0.0001 |
| | | | | P-γ15 vs P-γ2.5 | < 0.0001 |
| | | | | P-γ15 vs P-γ5 | < 0.0001 |
| | hiPS | P | 64 | P-γ2.5 vs P | < 0.0001 |
| | | P-γ2.5 | 63 | P-γ5 vs P | < 0.0001 |
| | | P-γ5 | 56 | P-γ5 vs P-γ2.5 | 0.84263 |
| | | P-γ10 | 66 | P-γ15 vs P | < 0.0001 |
| | | | | P-γ15 vs P-γ2.5 | 0.01309 |
| | | | | P-γ15 vs P-γ5 | 0.00104 |
| | 3T3-swiss | P | 132 | P-γ2.5 vs P | < 0.0001 |
| | | P-γ2.5 | 162 | P-γ5 vs P | 0.91497 |
| | | P-γ5 | 99 | P-γ5 vs P-γ2.5 | 1.05E-04 |
| | | P-γ10 | 130 | P-γ15 vs P | 0.66678 |
| | | | | P-γ15 vs P-γ2.5 | < 0.0001 |
| | | | | P-γ15 vs P-γ5 | 0.32326 |

| Related Figure | Cell | n | | P | |
|---|---|---|---|---|---|
| Fig. 6C | HeLa | A-γ2.5 | 20 | A-γ10 vs A-γ2.5 | < 0.0001 |
| | | A-γ10 | 8 | N-γ2.5 vs A-γ2.5 | 0.81758 |
| | | N-γ2.5 | 26 | N-γ2.5 vs A-γ10 | < 0.0001 |
| | | N-γ15 | 18 | N-γ15 vs A-γ2.5 | 0.15049 |
| | | | | N-γ15 vs A-γ10 | < 0.0001 |
| | | | | N-γ15 vs N-γ2.5 | 0.49135 |
| Fig. S2 | hiPS | A-γ2.5 | 13 | A-γ10 vs A-γ2.5 | 0.98869 |
| | | A-γ10 | 19 | N-γ2.5 vs A-γ2.5 | 0.80591 |
| | | N-γ2.5 | 10 | N-γ2.5 vs A-γ10 | 0.59225 |
| | | N-γ15 | 11 | N-γ15 vs A-γ2.5 | 0.99937 |
| | | | | N-γ15 vs A-γ10 | 0.97258 |
| | | | | N-γ15 vs N-γ2.5 | 0.87551 |
| Fig. 6C | 3T3-swiss | A-γ2.5 | 43 | A-γ10 vs A-γ2.5 | < 0.0001 |
| | | A-γ10 | 66 | N-γ2.5 vs A-γ2.5 | 0.12062 |
| | | N-γ2.5 | 86 | N-γ2.5 vs A-γ10 | 1.93E-03 |
| | | N-γ15 | 95 | N-γ15 vs A-γ2.5 | 0.05486 |
| | | | | N-γ15 vs A-γ10 | 0.00481 |
| | | | | N-γ15 vs N-γ2.5 | 0.98365 |

| Related Figure | Cell | n | | P |
|---|---|---|---|---|
| Fig. S3 | HeLa | A-γ10 | 8 | 0.00348 |
| | | Gel-γ6 | 65 | |
| | hiPS | A-γ10 | 19 | 0.52852 |
| | | Gel-γ6 | 30 | |
| | 3T3-swiss | A-γ10 | 66 | < 0.0001 |
| | | Gel-γ6 | 110 | |
